# Control of OPC proliferation and repopulation by the intellectual disability gene PAK1 under homeostatic and demyelinating conditions

**DOI:** 10.1101/2024.04.26.591153

**Authors:** Yan Wang, Bokyung Kim, Shuaishuai Gong, Joohyun Park, Meina Zhu, Evelyn M. Wong, Audrey Y. Park, Jonathan Chernoff, Fuzheng Guo

## Abstract

Appropriate proliferation and repopulation of oligodendrocyte progenitor cells (OPCs) determine successful (re)myelination in homeostatic and demyelinating brains. Activating mutations in p21-activated kinase 1 (PAK1) cause intellectual disability, neurodevelopmental abnormality, and white matter anomaly in children. It remains unclear if and how PAK1 regulates oligodendroglial development. Here, we report that PAK1 controls proliferation and regeneration of OPCs. Unlike differentiating oligodendrocytes, OPCs display high PAK1 activity which maintains them in a proliferative state by modulating PDGFRa-mediated mitogenic signaling. PAK1-deficient or kinase-inhibited OPCs reduce their proliferation capacity and population expansion. Mice carrying OPC-specific PAK1 deletion or kinase inhibition are populated with fewer OPCs in the homeostatic and demyelinated CNS than control mice. Together, our findings suggest that kinase-activating PAK1 mutations stall OPCs in a progenitor state, impacting timely oligodendroglial differentiation in the CNS of affected children and that PAK1 is a potential molecular target for replenishing OPCs in demyelinating lesions.

## Introduction

Local or intra-lesional proliferation and adequate OPC repopulation in demyelinating lesions determine effective myelin formation and repair (Fancy et al., 2011a) (Gallo and Deneen, 2014). Indeed, over one-third of chronic demyelinating lesions in the brain and spinal cord of multiple sclerosis (MS) exhibit significantly low numbers of OPCs (Boyd et al., 2013; Wolswijk, 2002). Greater depletion of OPCs than oligodendrocytes was also observed in some active MS lesions (Cui et al., 2013). OPCs become progressively depleted in chronic lesions where limited remyelination occurs in preclinical rodent models of demyelination (Mason et al., 2004). The “OPC depletion” concept is further supported by recent data demonstrating that impaired OPC repopulation is the major reason preventing successful myelin repair in larger animal models of demyelination (Cooper et al., 2023). Although extensive effort has been dedicated to defining how OPC terminal differentiation is regulated (Emery, 2010), it is equally important and necessary to study mechanisms underlying OPC proliferation and repopulation given OPC inadequacy or depletion in demyelinating lesions. Moreover, in the homeostatic brain of humans and rodents, OPCs actively proliferate and expand their population numbers during the neonatal and early postnatal ages (Huang et al., 2020) for subsequent terminal differentiation at the appropriate timings (Nishiyama et al., 2021). As such, impaired or uncontrolled OPC proliferation negatively impacts developmental myelination.

Group I p21-activated kinases (PAK1, PAK1, PAK3) are a family of serine/threonine kinases; they are inactivated by auto-inhibition which is dis-inhibited by binding to the 21 kDa GTP-binding proteins Cdc42 and Rac1 (Wang and Guo, 2022). PAK1 is an established oncogenic protein kinase and its hyperactivation leads to tumor cell proliferation, migration, invasion, and survival (Kanumuri et al., 2020; Ong et al., 2011; Shrestha et al., 2012). *De novo* mutations in group I *PAKs* are frequently reported in children of intellectual disability and neurodevelopmental disorders [see review (Wang and Guo, 2022)]. Notably, gain-of-function activating mutations in human *PAK1* lead to intellectual disability, macrocephaly, and white matter hyperintensity of T2-weighted brain MRI (Harms et al., 2018; Horn et al., 2019; Kernohan et al., 2019; Scorrano et al., 2023) which is indicative of white matter hypomyelination (Schiffmann and van der Knaap, 2009; Steenweg et al., 2010). Mechanistically, PAK1 regulates biological processes, for instance, cell growth, migration, and actin cytoskeleton remodeling, through its kinase activity-dependent [see review (Wang and Guo, 2022)] and -independent functions (Davidson et al., 2021; Higuchi et al., 2008; Wang et al., 2013). While PAK1’s function in neuronal development and function has been extensively scrutinized [see review (Nikolic, 2008), its role in oligodendroglial development is incompletely defined in the context of homeostasis and demyelination (Wang and Guo, 2022).

The current study aimed to define if and how PAK1 dysregulation affects early postnatal OPC development and to determine if PAK1 controls OPC repopulation in demyelinating lesions. To this end, we generated a series of transgenic mice, including *Pak1*-null, *Pak1*-floxed, and loxp-STOP-loxP-Pak1 inhibition peptide (*LSL-PID*) and crossed them with two different Cre driver lines to intervene PAK1 and its kinase activity. Using *in vitro* cultures and transgenic mice of OPC-specific PAK1 manipulations, we conclude that PAK1, acting through its kinase activity, maintained OPC in a proliferating progenitor state by modulating PDGFRa-mediated mitogenic signaling pathway and is required for OPC repopulation in demyelination lesions. Our findings provide new cellular and molecular insights into neurodevelopmental impairment in *PAK1*-mutated children and shed light on the therapeutic potential of PAK1-mediated OPC repopulation in MS lesions.

## Results

### PAK1 expression and its kinase activity in the brain and oligodendroglial lineage cells

Among group I PAKs, PAK1 is highly enriched in oligodendroglial lineage cells in the adult CNS (**Fig. S1A**) regardless of animal ages as demonstrated by scRNA-seq analysis (**Fig. S1B**). Oligodendroglia-enrichment of PAK1 transcripts was confirmed by qPCR assays of acutely magnetic-activated cell sorting (MACS)-purified brain cell populations in the early postnatal murine brain (**Fig. S1C-E**). We employed Western blot to assess the temporal dynamics of PAK1 protein and found that PAK1 was persistently expressed in the murine brain throughout postnatal development (**Fig. 1A**). Phosphorylation of PAK1 at Thr423 is a reliable marker for the kinase activation (Wang and Guo, 2022). In contrast to PAK1 expression, PAK1 kinase activity, indicated by p-PAK1(T423) levels, was evident during early postnatal ages by P14, a time-window when OPCs are actively proliferating and expanding their densities in the murine brain, and progressively diminished thereafter (**Fig. 1A**). *Pak1*-null transgenic mice were generated and authenticated PAK1 expression in our study (**Fig. 1B**). Fluorescent immunohistochemistry (IHC) showed that PAK1 was expressed in Sox10^+^ oligodendroglial lineage cells (**Fig. 1C**, left) which was completely abolished in *Pak1*-null mice (**Fig. 1C**, right). Oligodendroglial expression of PAK1 was further confirmed by *Pak1-eGFP* reporter mice we generated (**Fig. 1D**).

**Figure 1.**
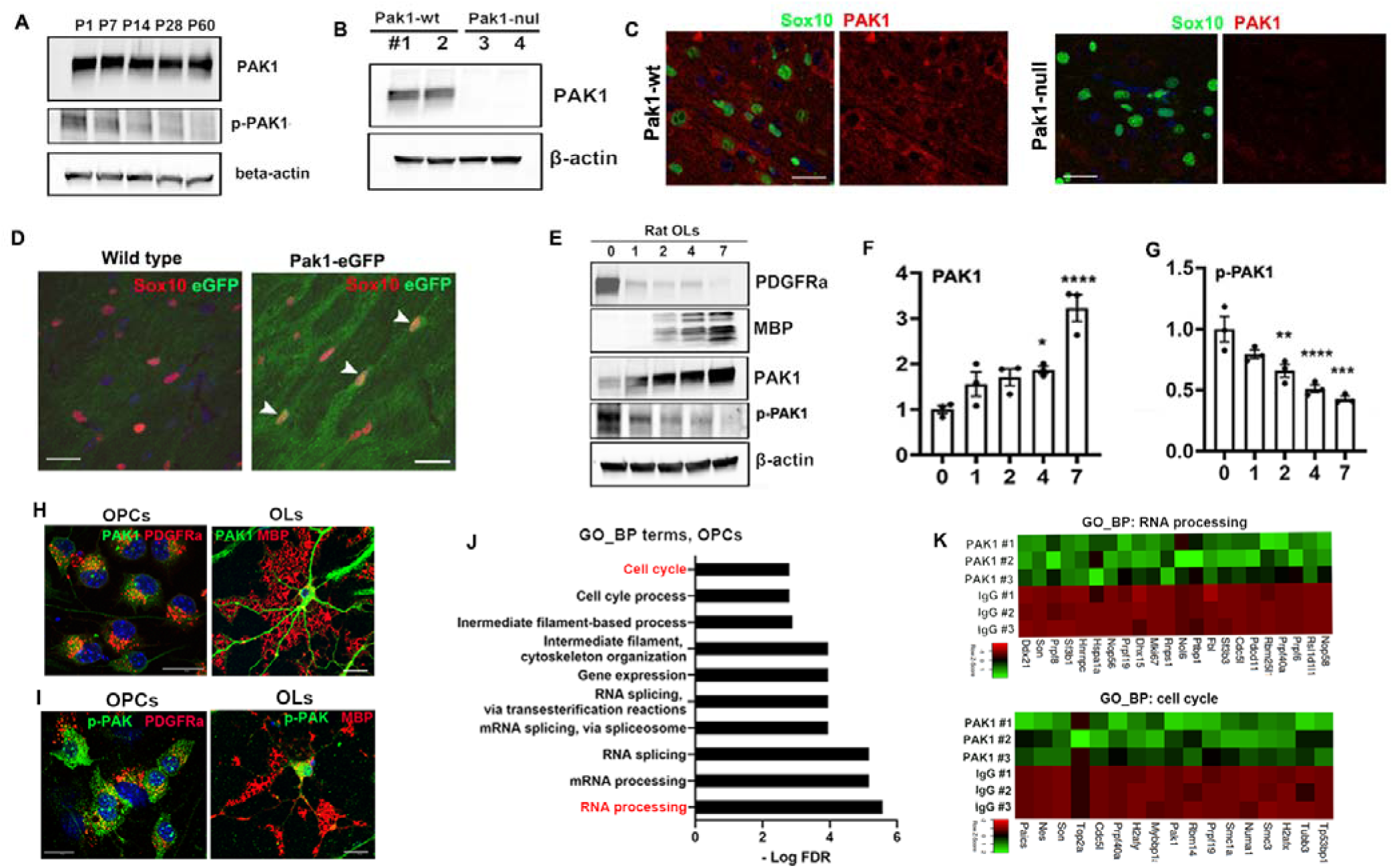
Expression and kinase activity of PAK1 in oligodendroglial lineage cells. **A**, Western blot (WB) assay of PAK1 and p-PAK (Thr423) in mouse brain, P, postnatal day. **B**, WB assay of PAK1 in P15 brain **C**, IHC of Sox10 and PAK1 on paraffin sections of P15 brain **D**, IHC of Sox10 and eGFP on frozen sections of P15 brain **E**, WB time course assay of PDGFRa, MBP, PAK1, and p-PAK1 (Thr423) in primary rat OPC (day 0), and differentiating OL at day 1, 2, 3, and 7. **F-G**, quantification of PAK1 and p-PAK1 protein levels (statistical parameters, see Table S1, hereafter) **H**, ICC of PAK1 with OPC marker PDGFRa at day 0 and OL marker MBP at day 4 of rat OLs **I,** ICC of p-PAK1 (Thr423) with PDGFRa and MBP. **J**, Gene ontology biological process (GO_BP) terms of PAK1’s interacting proteins in primary rat OPCs (see Table S2-4). K, heatmap of PAK1’s interacting proteins overrepresented in the GO_BP of RNA processing and cell cycle. Scale bars=10µm.

We next determined the temporal dynamics of PAK1 and its activity within oligodendroglial lineage cells. To this end, we sought primary rat OPC culture system. Western blot assay showed that PAK1 protein was low in PDGFRa-expressing OPCs and increased in differentiating MBP-expressing oligodendrocytes (OLs) (**Fig. 1E, F**). In line with the temporal dynamics in the brain (**Fig. 1A**), PAK1 kinase activity was high in OPCs and progressively decreased in differentiating OLs (**Fig. 1E, G**). Our conclusion was further corroborated by fluorescent immunocytochemistry (ICC) of PAK1 and p-PAK1(Thr423) (**Fig. 1H, I**). Taken together, our data demonstrate that, contrary to OLs, OPCs display high level kinase activity of PAK1. This finding suggests that PAK1 kinase activity plays a crucial role in controlling OPC behavior during early postnatal brain development.

### PAK1’s interactome in OPCs is primarily involved in cell cycle and RNA metabolism

PAK1 exerts biological functions primarily by phosphorylating and regulating its downstream target proteins, a process that requires PAK1 interaction with its target proteins. To profile PAK1’s interactome, we sought co-immunoprecipitation (IP) followed by unbiased LC-MS/MS protein identification (**Fig. S2A**). Two hundred forty-seven proteins (**Table S2**) were identified as potential binding partners in OPCs and differentiating OLs including PAK1 itself (**Fig. S2C**) and many other known targets (**Fig. S2D**). Surprisingly, the interactome was largely non-overlapped between OPCs and differentiating OLs; only 14.4% and 9.5% of binding proteins identified from OPCs and OLs were shared between the two populations (**Fig. S2B**). GO analysis demonstrated that PAK1 binding partners in OPCs were primarily involved in cell cycle, gene expression, and RNA metabolism (**Fig. 1J, K**, **Table S3**) whereas those in OLs were associated with actin cytoskeleton organization (**Fig. S2E, F, Table S4**). Given that OPCs are highly proliferative in the early postnatal brain of humans and rodents, our data suggest that PAK1 may control OPCs in a proliferating progenitor state through its kinase activity.

### PAK1 controls OPC proliferation and promotes its population expansion

To determine the functional significance of PAK1, we cultured primary OPCs from the neonatal brain of *Pak1*-null and wildtype (WT) mice by initially seeding 10,000 OPCs (**Fig. 2A**). After 4 days of expansion in the growth medium, *Pak1*-sufficient OPCs increased their population by ∼10 times whereas *Pak1*-deficient OPCs only by 3-4 times (**Fig. 2B**) suggesting that PAK1 is required for OPC population expansion. The percentage of Ki67^+^ cycling OPCs was significantly reduced in the *Pak1*-null group (**Fig. 2C**) which was corroborated by EdU pulse labeling (2 hours) of S-phase OPCs (**Fig. 2C**). We did not observe a marked difference in cleaved caspase 3 (CC3^+^) between *Pak1*-null and WT OPCs (data not shown). OPC displayed high levels of PAK1 kinase activity, therefore, we hypothesize that PAK1 controls OPC behavior through its kinase-dependent functions. To this end, we cultured *Pak1*-sufficient OPCs in the growth medium supplemented with or without PAK1 allosteric inhibitor IPA3 (**Fig. 2D**). Western blot showed that IPA3 inhibited PAK1 activity without affecting PAK1 protein expression (**Fig. 2E**). We found that PAK1 inhibition significantly impaired OPC population expansion (**Fig. 2F**) and proliferation capacity (**Fig. 2G**). In line with our *Pak1-null* OPC culture, no significant difference in CC3^+^ apoptotic OPCs was observed in IPA3 groups compared to the DMSO control group (**Fig. 2H**). Collectively, these data of genetic and pharmacological manipulations suggest that PAK1, acting through its kinase activity, is essential for OPC proliferation and population expansion.

**Figure 2.**
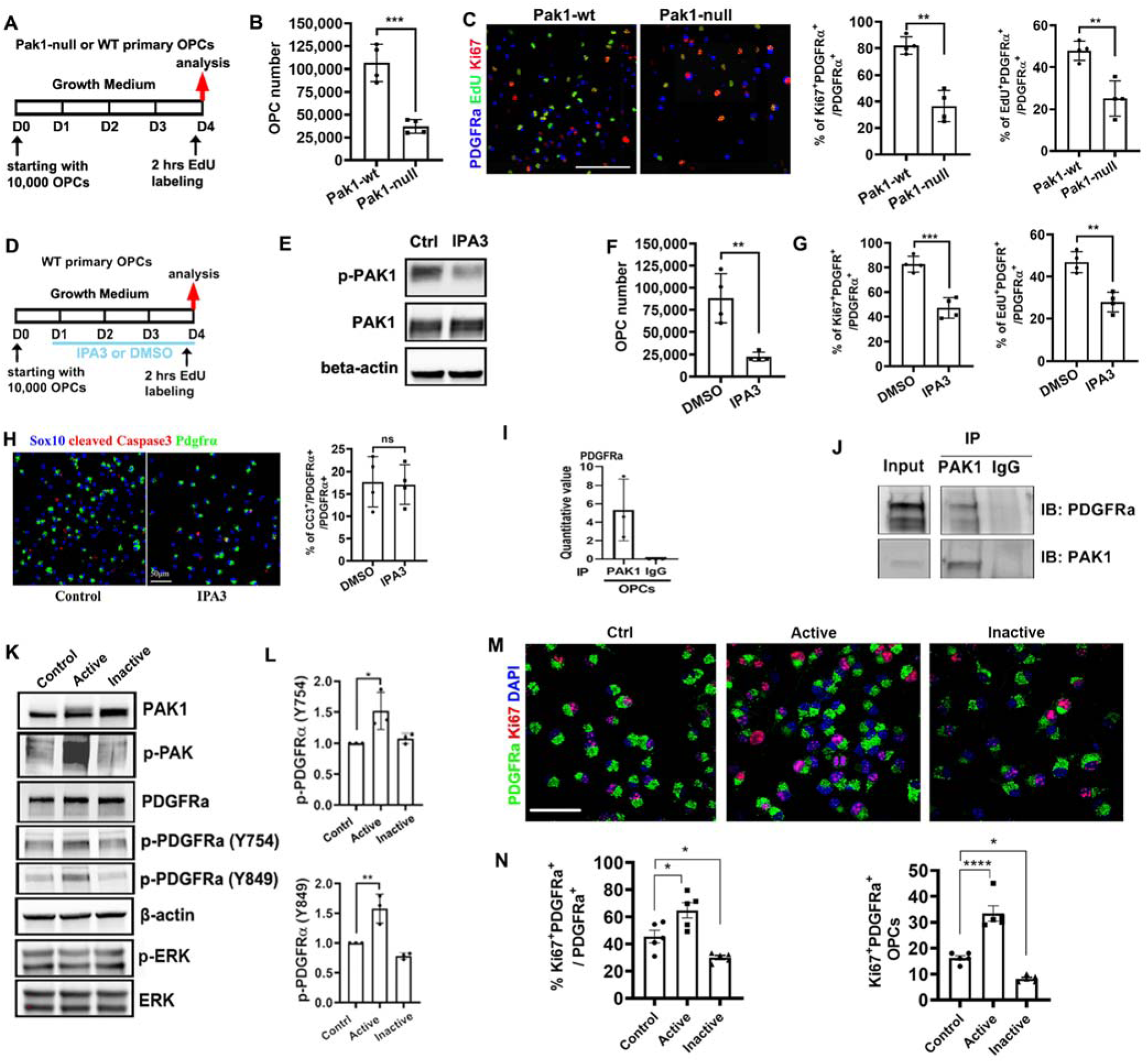
PAK1 controls OPC proliferation and population expansion through modulating PDGFRa signaling activity. **A,** experimental designs for panels B-C. **B**, total numbers of OPCs after 4 days of growth **C,** representative confocal images and quantification of EdU^+^ and Ki67^+^ proliferating OPCs **D**, experimental design for panel E-H **E**, WB assay of PAK1 and p-PAK1 (Thr423) at day 4 of OPC expansion **F**, total numbers of OPCs after 4 days of growth **G,** percentage of Ki67^+^ proliferating OPCs and EdU^+^ OPCs among total PDGFRa^+^ OPCs **H,** representative confocal image and quantification of cleaved caspase 3 (CC3), Sox10, and PDGFRa **I**, fold enrichment of PDGFRa in Co-IP by PAK1 antibody and IgG control **J**, Co-IP followed by immunoblot (IB) assay of PDGFRa-PAK1 interaction in OPCs **K**, WB assay of protein extracted from rat OPCs transfected with Ctrl, constitutive active, and dominant negative (inactive) PAK1 for 48 hours. **L**, quantification of p-PDGFRa at Y754 and Y849 versus total PDGFRa protein level **M**, representative confocal images of PDGFRa and Ki67 in rat OPCs at 48 hours after transfection **N**, quantification of percentage (left) and density (# per mm^2^, right) of Ki67^+^PDGFRa^+^ proliferating OPCs Scale bar: C, 100 µm; H, M, 50 µm.

### PAK1 modulates PDGFRa-mediated mitogenic signaling to maintain OPCs in a proliferative progenitor state

Our Co-IP and proteomics experiment showed that PAK1 interacted with PDGFRa (**Fig. 2I**), a key receptor that maintains OPCs in a proliferating progenitor state and prevents them from precocious differentiation (Cardona et al., 2021; Zheng et al., 2018). To verify the interaction of PAK1 with PDGFRa, we sought to employ Western blot assay after PAK1 Co-IP. Our results demonstrated that PDGFRa was detected in the IP elution of PAK1 antibody but absent from IgG control (**Fig. 2J**), suggesting that PAK1 interacts with PDGFRa in OPCs. To establish the functional regulation of PDGFRa-mediated signaling pathway by PAK1, constitutive active (PAK1-T423E) and dominant negative (PAK1-H83L/H86L/K299R) forms of PAK1 were expressed in primary rat OPCs cultured in the growth medium. Western blot demonstrated that PAK1-T423E expression, but not dominant negative form, significantly augmented PAK1 kinase activity (**Fig. 2K**). Expressing the kinase active PAK1 did not affect total PDGFRa expression but significantly increased PDGFRa activation, as demonstrated by elevated phosphorylation of PDGFRa at Tyr754 and Tyr849 (**Fig. 2K, L**), reliable readouts of PDGFRa signaling activation (Heldin and Lennartsson, 2013). Dominant negative PAK1 failed to modulate Tyr574/Tyr849 phosphorylation levels (**Fig. 2K, L**). In line with the IPA3 inhibition data, expression of the kinase active PAK1 increased whereas inactive PAK1 decreased the percentage of proliferating OPCs (**Fig. 2M-N**). Interestingly, we did not observe significant changes in MAPK-ERK1/2 pathway activation by active PAK1 expression (**Fig. 2K**) which has been previously reported in tumor cells (Coles and Shaw, 2002; Shrestha *et al*., 2012). Taken together, these data indicate that constitutive PAK1 activation, as reported in children with PAK1 activation mutations, maintains OPC in a proliferative state partially by modulating PDGFRa-mediated signaling pathway.

Given the progressive downregulation of PAK1 kinase activity in differentiating OLs (**Fig. 1E**), we speculated that PAK1 activity may act as a “brake” preventing OPC differentiation into OLs. To test this, we incubated primary rat OPCs with FRAX486 (PAK1 competitive inhibitor) and IPA3 (PAK1 steric inhibitor) in the differentiation medium (**Fig. S3A**) and determined the outcomes of inhibiting PAK1 activity on OPC differentiation. It was shown that IPA3 effectively inhibited PAK1 activity without affecting its expression (**Fig. S3B**) and that IPA3 did not affect OL viability up to the dose of 2.5µM as we assessed (**Fig. S3C**). We found that both IPA3 and FRAX486 treatment increased the expression of myelin genes *Mbp, Mog*, and *Opalin* at 3 days of differentiation with minimal effects on *Sox10* and *Pdgfra* compared to DMSO control (**Fig. S3D**). Fluorescent ICC (**Fig. S3E**) showed that the percentage of MBP^+^ cells among total Sox10^+^ cells was significantly increased upon inhibitor treatment (**Fig. S3F**). Sholl analysis demonstrated a more complex process network of IPA3 (or FRAX486)-treated oligodendrocytes than the control group (**Fig. S3G**). These data are in congruent with our hypothesis that PAK1 activity maintains OPCs in an undifferentiated state; its inhibition removes the “brake”, thus accelerating oligodendroglial lineage progression under differentiating conditions.

### PAK1 regulates OPC proliferation and population expansion *in vivo*

To determine the functional significance of PAK1 *in vivo*, *Pak1*-floxed mice were engineered for *Cre-loxP*-based conditional knockout (cKO) (**Fig. S4**). We generated *Sox10-Cre:Pak1*^fl/fl^ mice (**Fig. 3A**) in which *Sox10-Cre*-mediated *Pak1* gene disruption occurs initially in OPCs during CNS development. qPCR and Western blot assays showed a greater than 5-fold decrease in Pak1 mRNA (**Fig. 3B**) and protein (**Fig. 3C**) in the whole forebrain at P7, a time-window when OPCs are highly proliferative. The remarkable reduction of PAK1 expression in *Sox10-Cre:Pak1*^fl/fl^ whole forebrain indicates that Sox10-expressing oligodendroglial lineage cells are the major cell population expressing PAK1 among all brain cell types. We observed that OPC population density in the subcortical white matter (SCWM) was significantly reduced in *Sox10-Cre:Pak1*^fl/fl^ mice (**Fig. 3C, D**). The density of OPCs that are positive for phosphorylated histone H3 (PH3), a marker labeling cells undergoing mitosis (Kim et al., 2017), was significantly decreased in *Sox10-Cre:Pak1*^fl/fl^ mice (**Fig. 3C, E**), which was corroborated by decreased density of OPCs positive for Ki67, a nuclear marker labeling cells in the proliferating state (**Fig. 3F**). RT-qPCR confirmed the reduction in brain Ki67 transcripts (**Fig. 3F**). Together, these data suggest that PAK1 is essential for OPC proliferation and population expansion in the early postnatal brain.

**Figure 3.**
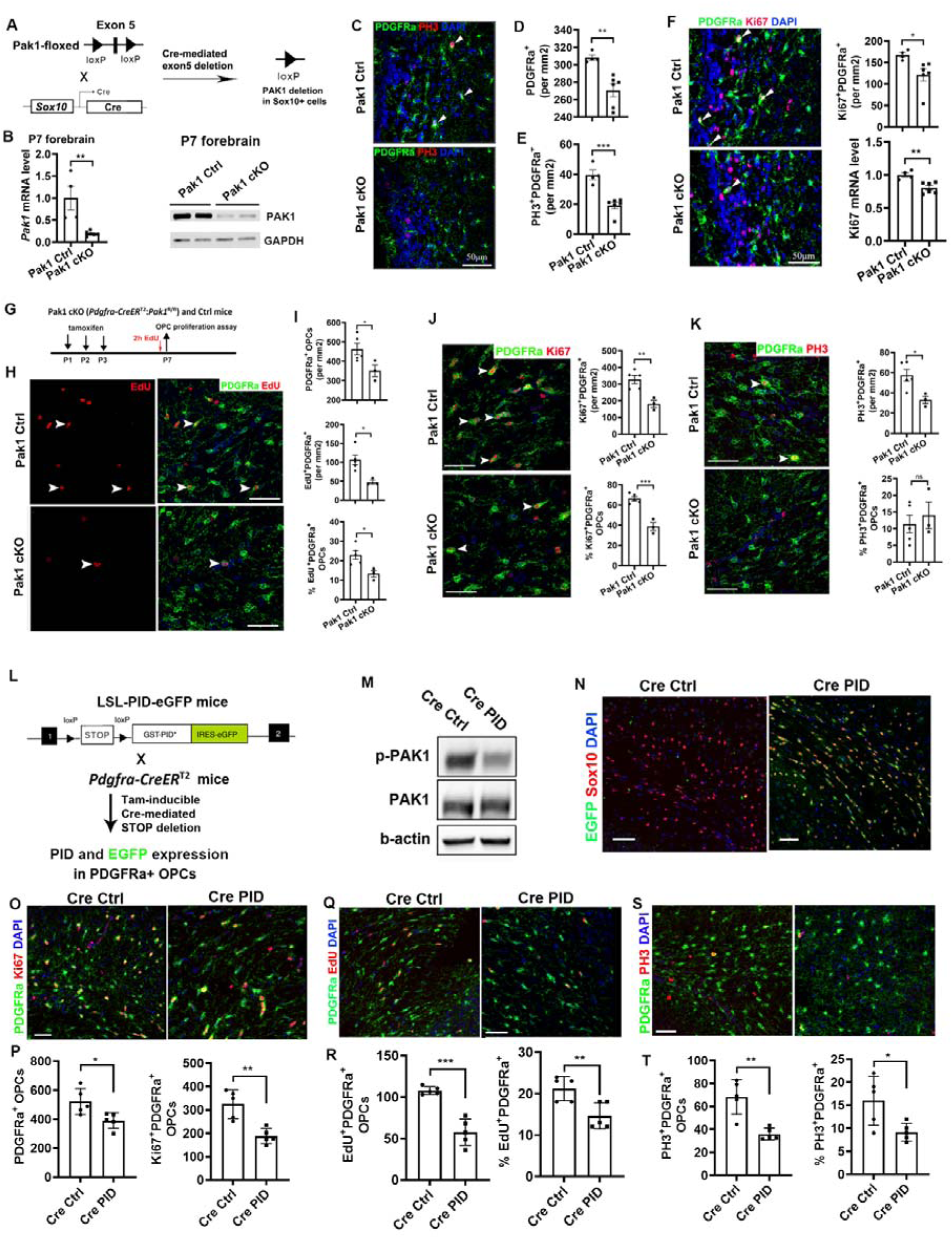
PAK1 controls OPC proliferation and expansion in vivo. **A**, experimental designs of Cre-loxP-mediated PAK1 conditional knockout (cKO) constitutively in Sox10-expressing OPCs. Pak1 cKO - *Sox10-Cre:Pak1*^fl/fl^, Pak1 Ctrl – *Sox10-Cre:Pak1*^+/+^, Panel C-F **B**, RT-qPCR (left) and WB assays of Pak1 mRNA and protein in P7 forebrain **C-E**, representative confocal images and quantification of PDGFRa^+^ OPCs and phosphor histone H3 (PH3)^+^PDGFRa^+^ mitotic OPCs in the brain subcortical white matter (SCWM). **G**, experimental designs of tamoxifen-inducible PAK1 cKO for Panel H-K. Pak1 cKO - *Pdgfra-CreER^T2^:Pak1*^fl/fl^, Pak1 Ctrl – *Pdgfra-CreER^T2^:Pak1*^+/+^ **H-I**, representative confocal image and quantification of PDGFRa^+^ OPCs, EdU^+^/ PDGFRa^+^ dividing OPCs, and percentage among total OPCs in the brain SCWM. **J**, representative confocal images, density of Ki67^+^PDGFRa^+^ proliferating OPCs, and percentage among total OPCs in the brain SCWM. **K**, representative confocal images, density of PH3^+^PDGFRa^+^ mitotic OPCs, and percentage among total OPCs in the brain SCWM. **L**, experimental designs of tamoxifen-inducible PAK inhibition (peptide inhibition domain, PID) for Panel M-T. Cre Ctrl - *Pdgfra-CreER^T2^*, Cre PID – *Pdgfra-CreER^T2^:LSL-PID.* Tamoxifen was i.p. administered to neonatal mice at P1, P2, P3. EdU was i.p. injected 2 hours prior to sacrifice at P7. PID expression is concomitant with EGFP expression in PDGFRa^+^ OPCs. **M**, WB assay of p-PAK1 (Thr423) and total PAK1 in the P7 brain. Beta-actin, internal protein loading control. **N**, EGFP expression in the SCWM of Cre:PID and Cre Ctrl mice. **O-P**, representative confocal images and densities (#/mm^2^) of PDGFRa^+^ total OPCs and Ki67^+^PDGFRa^+^ proliferating OPCs in the brain SCWM. **Q-R**, representative confocal images and density (#/mm^2^) and percentage of EdU^+^PDGFRa^+^ proliferating OPCs in the brain SCWM. **S-T**, representative confocal images and density (#/mm^2^) and percentage of PH3^+^PDGFRa^+^ proliferating OPCs in the brain SCWM. Scale bars=50 µm.

To strengthen our conclusion, we leveraged a time-controlled acute PAK1 cKO system by generating *Pdgfra-CreER^T2^:Pak1^fl/fl^*mice in which PAK1 was depleted at neonatal ages by tamoxifen administration (**Fig. 3G**). Consistent with *Sox10-Cre:Pak1*^fl/fl^ mice, *Pdgfra-CreER^T2^:Pak1^fl/fl^* mice displayed significantly reduced number and percentage of OPCs, EdU-labeling dividing OPCs (**Fig. 3H-I**), Ki67^+^ OPCs in cell cycles (**Fig. 3J**), and PH3^+^ OPCs undergoing mitosis (**Fig. 3K**).

### PAK1 kinase activity is essential for OPC proliferation and population expansion *in vivo*

To determine if PAK1 kinase activity plays a role in OPC proliferation and population expansion, neonatal mice were administered with IPA3 (**Fig. S5A**) to inhibit PAK1 kinase activity (**Fig. S5B**). We observed a significantly decreased number of total OPCs and Ki67^+^ OPCs in cell cycles (**Fig. S5C-E**) and PH3^+^ OPCs undergoing mitosis (**Fig. S5F-G**), suggesting that PAK1, acting through its kinase activity, regulates OPC population proliferation and expansion.

To define OPC-specific role PAK1 kinase activity, we employed *Cre-loxP* approaches to express PAK1 inhibitory domain (PID, amino acid residues 83-149 of PAK1), which binds to and inhibits the kinase domain of PAK1 (Chow et al., 2018; Wang and Guo, 2022). To achieve OPC specificity, we generated *Pdgfra-CreER*^T2^:*LSL-PID-eGFP* mice (Cre PID) (**Fig. 3L**). Our prior data showed that PAK activity was effectively inhibited by PID expression in eGFP^+^ cells (Chow *et al*., 2018). In the brain, the kinase activity of PAK1, but not its expression, was remarkably reduced in Cre PID mice versus Cre Ctrl mice (**Fig. 3M**). Fluorescent IHC further confirmed the cellular specificity of PAK inhibition, as almost all PAK-inhibited/eGFP-expressing cells were Sox10^+^ cells in the brain of *Pdgfra-CreER*^T2^:*LSL-PID-eGFP* mice (**Fig. 3N**). Built on this effective system, we observed that the densities of total OPCs, Ki67^+^ OPCs in cell cycles, EdU^+^ dividing OPCs, and PH3^+^ OPCs undergoing mitosis were all significantly decreased in *Pdgfra-CreER*^T2^:*LSL-PID-eGFP* mice compared to controls (**Fig. 3O-T**).

Collectively, our data suggest that PAK1-regulated OPC proliferation and population expansion are, at least in part, through its kinase activity.

### PAK1 is required for OPC proliferation and lesional repopulation in response to demyelination

OPC recruitment (repopulation) and intra-lesional proliferation in demyelinating areas are prerequisites for myelin repair and functional recovery. OPC depletion and inadequate repopulation are hallmarks for many chronically demyelinating plaques in multiple sclerosis. To determine if PAK1-regulated OPC behavior is preserved in demyelinating disorders, we leveraged the well-established focal demyelination mouse model of lysolecithin in which both OLs and OPCs are depleted in the lesional area. P60 *Pdgfra-CreER*^T2^:*Pak1*^fl/fl^ and *Pdgfra-CreER*^T2^ control mice were administered with tamoxifen followed by stereotaxic injection of lysolecithin into the corpus callosum 2 weeks after (**Fig. 4A**). The treated mice were killed for histological analysis at 5 days post-lesioning (5 dpl) when peak OPC recruitment and proliferation occur in the model (Fancy et al., 2009; Fancy et al., 2011b). Neural red (NR) dye was used to objectively identify lysolecithin-elicited lesion areas (Baydyuk et al., 2019) in our experimental designs. PAK1 deletion resulted in significant reduction of OPC densities in NR-labeled demyelination lesions of *Pdgfra-CreER*^T2^:*Pak1*^fl/fl^ compared to control mice (**Fig. 4B, C**). Furthermore, EdU pulse labeling showed a significant decrease in the number and percentage of actively dividing OPCs (**Fig. 4B, D**). Thus, PAK1 is required for OPC recruitment and intra-lesional proliferation in response to demyelination injury.

**Figure 4.**
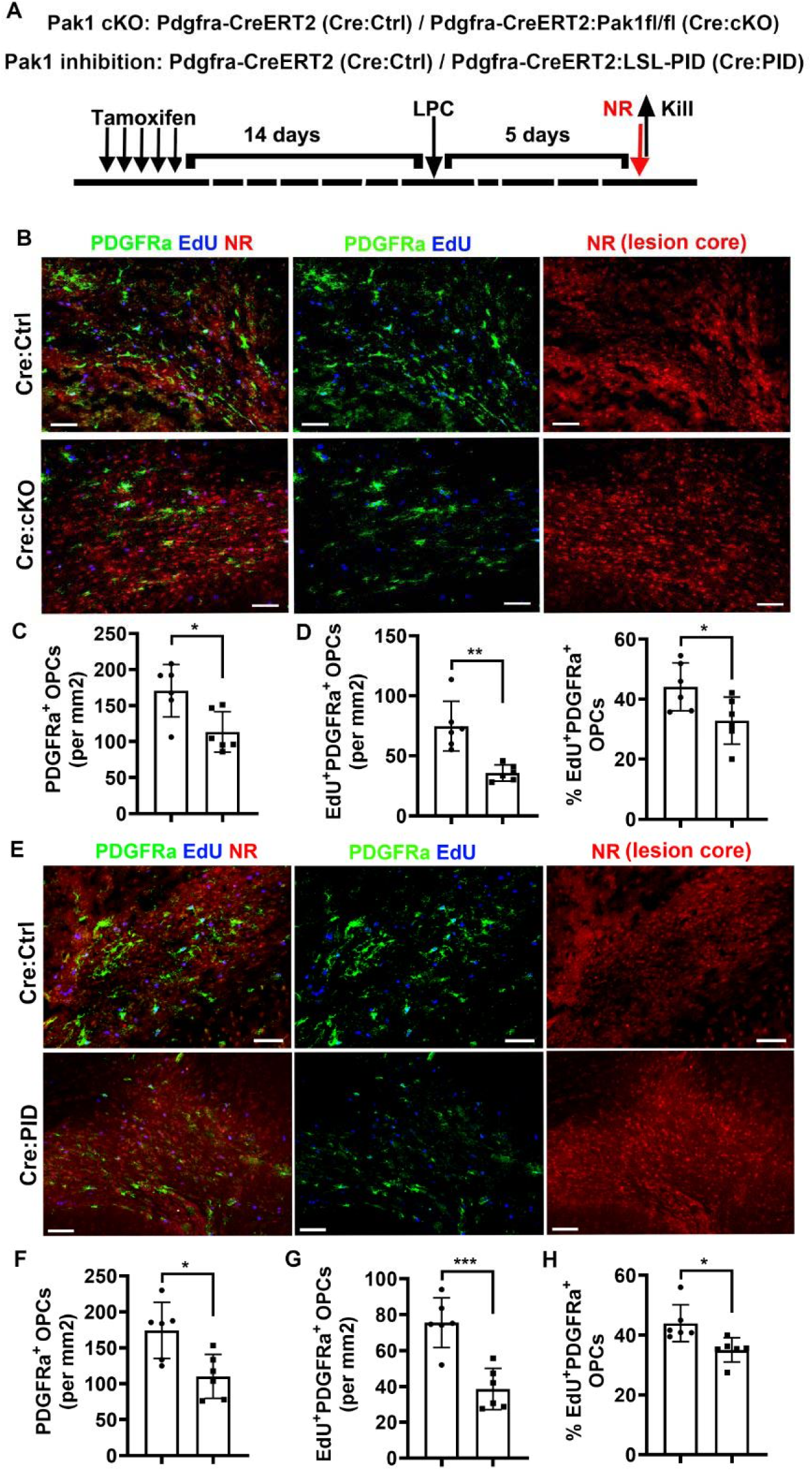
PAK1 controls OPC proliferation and lesion repopulation after demyelination through its kinase activity. **A**, experimental designs for panel B-G. P60 transgenic mice carrying Pak1 cKO, PID, and respective Cre Ctrl were i.p. injected with 5-day tamoxifen. After 2 weeks clearance time, treated mice were induced focal demyelination by lysolecithin (lysophosphatidylcholine LPC) stereotaxic injection into the corpus callosum and killed at 5 days post-lesioning (dpl), a time point of active OPC proliferation and lesion recruitment. Neural red (NR) was i.p. injected 2 hours prior to sacrifice to track lesions. **B-D**. confocal images (**B**) and quantification of PDGFRa^+^ OPCs (**C**), EdU^+^PDGFRa^+^ dividing OPCs and percentage (**D**) in lesion cores of *Pdgfra-CreER^T2^:Pak1*^fl/fl^ (Pak1 cKO) versus *Pdgfra-CreER*^T2^ control mice. **E-H**. confocal images (**E**) and quantification of PDGFRa^+^ OPCs (**F**), EdU^+^PDGFRa^+^ dividing OPCs (**G**) and percentage (**H**) in lesion cores of *Pdgfra-CreER^T2^:LSL-PID* (Pak inhibition) versus *Pdgfra-CreER*^T2^ control mice.

### PAK1-regulated OPC lesional repopulation is mediated by its kinase activity

To determine if PAK1 activity plays a role, *Pdgfra-CreER*^T2^:*LSL-PID-eGFP* and control mice were subjected to the same experimental perturbations as *Pdgfra-CreER*^T2^:*Pak1*^fl/fl^ mice. PAK activity inhibition resulted in a significant decrease in total PDGFRa^+^ OPCs in NR-positive lesioned areas (**Fig. 4E, F**). EdU pulse labeling experiments demonstrated that the density (**Fig. 4G**) and percentage of Edu-incorporated (**Fig. 4H**) OPCs in NR-positive lesion areas of *Pdgfra-CreER*^T2^:*LSL-PID-eGFP* mice compared to those in *Pdgfra-CreER*^T2^ Ctrl mice. Taken together, our genetic evidence demonstrated that PAK1’s kinase activity is essential for OPC proliferation and repopulation in demyelination lesions in response to white matter injury.

## Discussion

Mutations of the auto-inhibitory domain of PAK1 result in constitutive kinase activation in the brains of affected children and cause neurodevelopmental disturbance, macrocephaly, and intellectual disability. By employing a series of transgenic mice targeting PAK1 and its kinase activity in combination with *in vitro* molecular manipulations and pharmacological inhibition, the current study provided convincing evidence that PAK1, acting through its kinase activity, maintains OPCs in a proliferating progenitor state by modulating PDGFRa-mediated mitogenic signaling pathway and is required for OPC population expansion. As such, *Pak1*-deficient or kinase-inhibited OPCs decrease their proliferation capacity and population numbers both *in vitro* and *in vivo*. It is well-established that the mitogenic pathway mediated by PDGFRa is required for OPC migration and proliferation (Zheng *et al*., 2018). This study provides new evidence linking PAK1 function with PDGFRa-mediated mitogenic signaling in OPCs, as we observed that constitutive kinase activation of PAK1 significantly enhanced PDGFRa activation and promoted OPC proliferation (Fig. 2K-N). Our findings suggest that kinase activating mutations in human PAK1 may result in prolonged OPC proliferation and perturb the timing of brain myelination, potentially contributing to neurodevelopmental defects in affected children (Harms *et al*., 2018; Horn *et al*., 2019; Kernohan *et al*., 2019; Scorrano *et al*., 2023). PDGFRa belongs to the receptor tyrosine kinase (RTK) family that are activated by growth factors (Chen et al., 2013). A previous study showed that PAK1 kinase is stimulated by PDGFRa upon growth factor binding in cancer cells (Galisteo et al., 1996). The data of the current study demonstrated that PAK1 kinase function stimulates PDGFRa activation in OPCs, suggesting a reciprocal regulation between PAK1 and PDGFRa. Our finding is also in agreement with previous observation that *Pdgfra* expression is significantly downregulated by PAK1 KO in mouse embryonic fibroblasts (Sanchez-Solana et al., 2012). It remains unclear how PAK1 modulates OPC PDGFRa activation. Given that PAK1 regulates its downstream targets by phosphorylating Ser/Thr residues, we postulate that PDGFRa-regulation by PAK1 may occur in an indirect way mediated by yet-to-be-identified proteins in OPCs (Wang and Guo, 2022).

PAK activity was evident in the neonatal murine brain where active OPC proliferation occurs and was sharply downregulated after P14 (Fig. 1A). Moreover, the activity downregulation is temporally associated with OPC terminal differentiation during oligodendroglial lineage progression and maturation (Fig. 1E). The temporal correlation suggests that PAK1 kinase activity downregulation is a prerequisite or permissive for subsequent OPC differentiation into oligodendrocytes. We found that dampening PAK activity by two small inhibitors of distinct inhibitory mechanisms accelerated myelin gene expression and OPC terminal differentiation into oligodendrocytes only under the differentiating conditions (Fig. S3). Our observation is consistent with a recent preprint showing that constitutively active PAK1 expression decreased myelin membrane formation *in vitro* under the differentiating conditions – an important metrics of oligodendroglial morphological differentiation (Baudouin et al., 2023). The findings from Baudouin *et al* and the current study suggest that PAK1 activity may act as a molecular brake limiting oligodendrocyte lineage progression and differentiation in rodents. In contrast, a previous study reported that PAK1 was a positive regulator for oligodendroglial morphological differentiation in primary rodent OPC cultures (Brown et al., 2021) where PAK1 or its activity was experimentally perturbed in the OPC stages. It is plausible that dampening OPC levels of PAK1 or its kinase activity interferes with OPC proliferation/population, potentially impacting OPC differentiation capacity. Taken together, these data prompted us to hypothesize a dual-modal role of PAK1 in oligodendroglial lineage development: it promotes OPC proliferation and population expansion and later inhibits oligodendroglial differentiation. To define the *in vivo* role of PAK1 and its kinase activity in OPC terminal differentiation into oligodendrocytes, further studies are needed to generate transgenic mice carrying oligodendrocyte-specific PAK1 cKO or PID expression in which OPC proliferation and population are unperturbed.

Given the shared autoinhibition mechanisms among group I PAKs (Wang and Guo, 2022), pharmacological inhibition of PAK1 activity might also perturb PAK2 or PAK3 activity in our and others’ studies (Baudouin *et al*., 2023; Brown *et al*., 2021). Similarly, PID expression inhibits the activity of not only PAK1 but also PAK2 or PAK3. Our *Pak1* KO data suggest that PAK1 regulation of OPC proliferation and population expansion is not fully compensated by PAK2 or PAK3. PAK1 is relatively enriched in oligodendroglial lineage whereas PAK3 expression is highest in neurons in the young adult and aged brain (Fig. S1A-B). In the early postnatal brain, we found that PAK1 and PAK3 were enriched in oligodendroglia whereas PAK2 was ubiquitously expressed in brain cells (Fig. S1E). Functionally, PAK1 KO and PAK3 KO mice are viable whereas PAK2 KO mice die during early embryonic development due to vascular malformation. PAK3 also appears to regulate OPC proliferation, as global PAK3 KO mice display transient delay in OPC proliferation and terminal differentiation in restricted brain white matter tracts (Maglorius Renkilaraj et al., 2017). Clinically, loss-of-function mutations of human *PAK2* gene result in autism (Wang et al., 2018) where an accelerated white matter maturation and hypermyelination are observed in the brain compared with healthy controls (Ben Bashat et al., 2007), suggesting that PAK2 may inhibit OPC terminal differentiation. It remains enigmatic if PAK1 and PAK2 (or PAK3) are synergistic or redundant in regulating OPC terminal differentiation and/or myelination in the brain. To provide novel insights into the enigma, our group is generating transgenic mice carrying oligodendrocyte-specific single depletion of PAK1 or PAK2 and double deletion of PAK1/PAK2.

Another interesting finding of this study is that PAK1, acting through its kinase activity, is necessary for OPC repopulation in lesions after focal demyelination (Fig. 4). This finding is significant in that OPC depletion was observed in a large proportion of MS chronic lesions. In addition to differentiation blockade, OPC population insufficiency has been proposed as a reason for incomplete or failed myelin repair. Indeed, newly regenerated oligodendrocytes within MS lesions were only present in significant numbers with high OPC densities (Wolswijk, 2002). Spontaneous remyelination is extensive in some MS lesions (Patrikios et al., 2006) with sufficient OPC proliferation and OPC repopulation. We reported for the first time that PAK1 depletion or kinase inhibition significantly impaired OPC proliferation and repopulation in focal demyelinating lesions. Our observation is in line with previous data showing that the application of FTY720 (Fingolimod), a potent stimulator of PAK1 activity (Bokoch et al., 1998; Egom et al., 2010; Liu et al., 2011; Roig et al., 2001), to cuprizone-fed mice promoted OPC proliferation and repopulation in demyelinating lesions (Kim et al., 2011). Our finding indicates that PAK activity may be a therapeutic target by which could be harnessed to promote OPC proliferation and repopulation for successful myelin repair in certain MS chronic lesions with OPC depletion.

### Limitations of the study

First, the transgenic mice engineered in the current study were powerful in deciphering the role of PAK1 in OPC proliferation and population expansion. Yet they were ineffective in studying the role of PAK1 in oligodendrocyte differentiation and myelination which will be negatively impacted by defective OPC proliferation and population expansion in the transgenic lines. In this regard, transgenic mice carrying oligodendrocyte (but not OPC)-specific PAK1 cKO or conditional inhibition will provide new insights into the important yet controversial topic. Given the inverse correlation between the levels of PAK1 protein expression and its kinase activity in mature oligodendrocytes, it is also interesting to investigate the kinase-independent roles of PAK1 in regulating oligodendrocyte functions in the adult CNS. Second, the current manuscript lacks detailed molecular mechanistic insights into how PAK1 modulates PDGFRa signaling activity in OPCs. This topic warrants future studies using immortalized OPC cell lines (such as Oli-Neu cells) which, unlike primary rodent OPCs, are resistant to various molecular manipulations. Lastly, transgenic mice carrying *Pak1* gain-of-function mutations equivalent to human PAK1 gene are not available in the current study to mimic neurodevelopmental disorders reported in PAK1-mutated children.

## Materials and Methods

### Generation of transgenic mice

#### Pak1-eGFP mice

*Pak1-eGFP* transgenic line was obtained from the Mutant Mouse Resource & Research Centers [MMRRC # 010583-UCD, Tg(Pak1-EGFP)EJ8Gsat/Mmucd] which includes the enhanced green fluorescent protein (eGFP) coding sequence, followed by a polyadenylation signal, inserted into the mouse genomic bacterial artificial chromosome (BAC) RP23-126P9 at the start codon of Pak1 gene. This ensures that the eGFP reporter’s expression is controlled by the regulatory elements of Pak1 gene. The genotype can be identified using specific primers: Forward 5’-GGAGATGGATGAATGGGACTGAA-3’, and Reverse 5’-GGTCGGGGTAGCGGCTGAA-3’, producing a 550bp fragment for the transgene. *Pak1-eGFP* mice were generated by the Mouse Biology Program (MBP) at UC Davis.

#### Pak1-null mice

*Pak1-null* mice (B6.129S2-*Pak1^tm1Cher^*/Mmnc) were obtained from the Mutant Mouse Resource & Research Centers (MMRRC, #031838-UNC). In the *Pak1-null* mice, the Neo cassette was inserted into exon 4, generating a premature STOP codon in exon 5 (which encodes a p21-binding domain) and causing premature truncation of protein. Genotype primer sequences: Common-Forward: 5’-GCCCTTCACAGGAGCTTAATGA-3’; WT-reverse: 5’-GAAAGGACTGAATCTAATAGC-3’; Neo-reverse: 5’-CATTTGTCACGTCCTGCACGA-3’. Agarose gel electrophoresis bands: mutant 360bp WT 240bp.

#### *Pak1*-floxed mice

A *Pak1*-floxed transgenic line was generated through a targeting strategy that involved the insertion of two loxP sites flanking the 38-bp exon 5 of *Pak1* gene (Supplementary Figure 4A). This manipulation was designed to allow the conditional deletion of exon 5. The WT PAK1 protein consists of 544 amino acids with an approximate molecular weight of 65 kDa. The engineered deletion of exon 5 introduces a premature stop codon (TGA) at the 18th nucleotide of exon 6, resulting in a truncated PAK1 protein comprising 151 amino acids with an estimated weight of 18 kDa.

To facilitate the identification of the floxed alleles (Tm1c, fl/fl), a pair of primers, F1 (5’-GGACAACCTTGGGTATATTCCTCAG-3’) and R1 (5’-GAGAGAAGTTAAGTAATTTGCCCAGC-3’), was designed for PCR amplification of genomic DNA. PCR analysis of genomic DNA from transgenic mice yielded the expected band sizes: 445 bp for the WT alleles (+/+), 567 bp for the floxed alleles (fl/fl), and a combination of both for the heterozygous (fl/+) condition (Supplementary Figure 4B). Additionally, primers F2 (5’-CGCTTGCTTCAAACATCAAATA-3’) and R2 (5’-GGTAGCATCGTCATCATCATCT-3’) were designed to detect the presence of *Pak1* exon 5-deleted alleles (-/-) in cDNA synthesized from mRNA isolated from Cre-positive cells. The conditional deletion of exon 5 was confirmed by PCR amplification of cDNA prepared from brain oligodendroglial cells. These cells were isolated using magnetic-assisted cell sorting (MACS) with O4 antibody-conjugated magnetic microbeads from postnatal day 7 (P7) mice carrying *Pa*k1^fl/fl^ and *Sox10-Cre:Pak1*^fl/fl^. The PCR results displayed a band size of 240 bp for the floxed allele prior to Cre-mediated recombination and a band size of 202 bp post-exon 5 deletion, validating the expected genetic alteration in Cre-expressing cells (Supplementary Figure 4C). Tamoxifen was dissolved in a mixture of ethanol and sunflower seed oil (1:9, v/v) at the concentration of 30 mg/ml administered at a dose of 75mg/kg body weight to *Pdgfra-CreER^T2^* mice as indicated in each figure.

#### Pak inhibitor domain (PID) transgenic mice

PID, a peptide derived from the minimal autoinhibitory domain from Pak1 (minimally, corresponding to residues 83-149), was generated and characterized in our previous study (Chow *et al*., 2018). In the PID transgenic mouse, the transcription of *GST-PID* is blocked by the presence of upstream “lox-stop-lox” (LSL) sequences. The LSL-PID mice were crossed with Pdgfra-CreERT2 mice. Upon exposure to Cre recombinase in the nucleus, the “stop” sequences between the *loxP* sites are excised, leading to expression of the GST-PID transgene. In the meanwhile, the IRES (internal ribosome entry site) allows eGFP expression in Cre positive PID-expressing cells. For *Pdgfr*α*-CreERT2:PID* and littermate controls, tamoxifen was administered subcutaneously to neonatal pups on P1, P2, and P3 daily at a dose of 75 mg/kg body weight once a day and mice were killed at P7 for OPC proliferation assay.

#### OPC primary culture and proliferation assay

Primary mixed glial cells were isolated from the cerebral cortices of P0-P2 mice or rats, following our established protocols (Lang et al., 2013; Wang et al., 2021), and cultured in T75 flasks coated with poly-D-lysine (PDL). The culture medium consisted of high-glucose DMEM, supplemented with 10% heat-inactivated fetal bovine serum (FBS) and 1% Penicillin/Streptomycin (P/S). The medium was refreshed every two days until the astrocyte layer reached confluency. To isolate microglia, the flasks were subjected to orbital shaking at 37°C and 200 rpm for 1 hour (Orbital shaker, Cat# C491, Hanchen), followed by a PBS wash. Subsequently, 20 mL of DMEM with 10% FBS was added to each flask. A further 6-hour shake under the same conditions facilitated the collection of OPCs. The OPCs were then seeded on PDL-coated plates using serum-free complete growth medium (CGM). The CGM was formulated from 30% B104 neuroblastoma-conditioned medium and 70% N1 supplement (DMEM enriched with 5 mg/mL insulin, 50 mg/mL apo-transferrin, 100 µM putrescine, 30 nM sodium selenite, 20 nM progesterone, 5 ng/mL FGF (Peprotech), 4 ng/mL PDGF-AA (Peprotech), 50 µM Forskolin (Peprotech), and GlutaMAX (Thermo Fisher). For differentiation, OPCs were transitioned to differentiation medium (DM) composed of F12/high-glucose DMEM, supplemented with 12.5 mg/mL insulin, 100 µM Putrescine, 24 nM Sodium selenite, 10 nM Progesterone, 10 ng/mL Biotin, 50 mg/mL Transferrin, 30 ng/mL 3,3’,5-Triiodo-L-thyronine, 40 ng/mL L-Thyroxine, GlutaMAX (Thermo Fisher), and P/S (Thermo Fisher) (all components from Sigma, unless noted).

#### Protein extraction and western blotting assay

Tissues or cells were lysed using N-PER Neuronal Protein Extraction Reagent (Thermo Fisher), supplemented with both a protease and phosphatase inhibitor cocktail (Thermo Fisher) and phenylmethylsulfonyl fluoride (PMSF; Cell Signaling Technology). The lysates were incubated on ice for 10 minutes, followed by centrifugation at 10,000 x g for 10 minutes at 4°C. Protein concentrations in each sample were then determined using the BCA Protein Assay Kit (Thermo Fisher Scientific). Equal amounts of cell lysates (30 µg) from each experimental condition were separated on either Any kD Mini-PROTEAN TGX Precast Gels or 7.5% Mini-PROTEAN TGX Precast Gels (BIO-RAD), according to the protein’s size. Proteins were subsequently transferred to 0.2 µm nitrocellulose membranes (BIO-RAD) using the Trans-Blot Turbo Transfer System (BIO-RAD). Membranes were blocked with 5% bovine serum albumin (BSA; Cell Signaling) for 1 hour at room temperature, then incubated overnight at 4°C with primary antibodies, followed by appropriate horseradish peroxidase (HRP)-conjugated secondary antibodies. The targeted proteins were visualized using Western Lightning Plus ECL (Perkin Elmer), and protein levels were quantified with NIH ImageJ software. Primary antibodies used were: PAK1 (1:1000, #2602, Cell Signaling Technology), P-PAK1(Ser144) (1:1000, #2606, Cell Signaling Technology), PDGFRα (1:1000, #AF1062, NOVUS), P-PDGFRα(Y754) (1:1000, #ab5460, Abcam), P-PDGFRα(Y849) (1:1000, #ab79318, Abcam), MBP (1:1000, #NB600-717, Novus), ERK1/2 (1:1000, #9102, Cell Signaling Technology), P-ERK1/2 (Thr202/Tyr204) (1:1000, #9101, Cell Signaling Technology), β-actin (1:1000, #4967, Cell Signaling Technology), and GAPDH (1:1000, #2118, Cell Signaling Technology). HRP-conjugated secondary antibodies (1:3000) are all from Thermo Fisher Scientific.

#### Co-Immunoprecipitation (Co-IP) and LC-MS/MS Analysis

Co-IP/LC-MS/MS assays were conducted utilizing the Pierce Classic Magnetic IP/Co-IP Kit (Thermo Fisher) in accordance with the manufacturer’s protocols. Protein lysates (1 mg) underwent overnight incubation with 10 µg of either PAK1 primary antibody (#2602, Cell Signaling Technology) or Rabbit IgG isotype control (#2729, Cell Signaling Technology) at 4°C. Subsequently, Pierce Protein A/G Magnetic Beads were washed three times using Pierce IP Lysis/Wash Buffer, followed by four times wash with 200 µL of 50 mM ammonium bicarbonate (AMBIC), shaking for 20 minutes at 4°C. Then, 2.5 µg of Trypsin Gold (Mass Spectrometry Grade, V528A, Promega) was introduced to the beads for overnight digestion at a shaking speed of 800 rpm at room temperature. Following digestion, peptide extracts were concentrated via vacuum centrifugation. An aliquot of the concentrated extract underwent fluorometric peptide quantification using the Thermo Scientific Pierce kit. The UC Davis Proteomics Core facilitated the LC-MS/MS analysis. Each analysis was performed with 1 µg of sample, as determined by the fluorometric peptide assay. These samples were analyzed on a Thermo Scientific Q Exactive Plus Orbitrap Mass Spectrometer, coupled with a Proxeon Easy-nLC II HPLC and Proxeon nanospray source. Proteome Discoverer (Thermo Scientific) was employed for tandem mass spectra extraction and charge state deconvolution. The X! Tandem software (The GPM, thegpm.org; version X! Tandem Alanine 2017.2.1.4) was used for analyzing all MS/MS samples, and Scaffold (version 4.8.4, Proteome Software Inc., Portland, OR) verified MS/MS-based peptide and protein identifications.

#### Co-Immunoprecipitation (Co-IP) and western blotting assay

Co-IP was carried out using the Pierce Crosslink Magnetic IP/Co-IP Kit (Thermo Fisher), adhering to the provided instructions. PAK1 primary antibody (#2602, Cell Signaling Technology) or Rabbit IgG isotype control (#2729, Cell Signaling Technology) (10 µg) were covalently cross-linked to 25 µL of Protein A/G Magnetic Beads (Thermo Fisher). Protein extraction was performed using Pierce IP Lysis/Wash Buffer (Thermo Fisher), supplemented with PMSF (Cell Signaling Technology) and a protease and phosphatase inhibitor cocktail (Thermo Fisher). A fraction of each sample was reserved as input. Equal amounts of protein extracts (1 mg) were incubated with the antibody- or IgG-cross-linked Protein A/G magnetic beads overnight at 4°C. Post-incubation, the beads were washed to eliminate unbound material, and proteins were eluted using a low-pH elution buffer, which facilitates the dissociation of the antigen from the antibody-bead complex. The eluates were neutralized with Neutralization Buffer and prepared for SDS-PAGE and Western blotting by adding Lane Marker Sample Buffer containing β-mercaptoethanol.

#### RNA extraction and quantitative real-time PCR (qRT-PCR)

RNA was isolated using the RNeasy Lipid Tissue Mini Kit (QIAGEN), and genomic DNA contamination was eliminated with the RNase-Free DNase Set (QIAGEN). The concentration of RNA was determined using a Nanodrop 2000 Spectrophotometer (Thermo Fisher Scientific). Subsequently, cDNA was synthesized employing the QIAGEN Omniscript RT Kit (QIAGEN). The RT-qPCR analyses were conducted with the QuantiTect SYBR Green PCR Kit (QIAGEN) on an Agilent MP3005P thermocycler. mRNA expression levels of target genes in each sample were normalized to the internal control gene Hsp90. Fold changes in gene expression were calculated using the 2^(-ΔCt) method, where ΔCt represents the difference between the cycle threshold (Ct) of Hsp90 and the Ct of the target genes. The expression levels of genes in control samples were set to a baseline value of 1 for comparative purposes. Primer sequences are as follows.

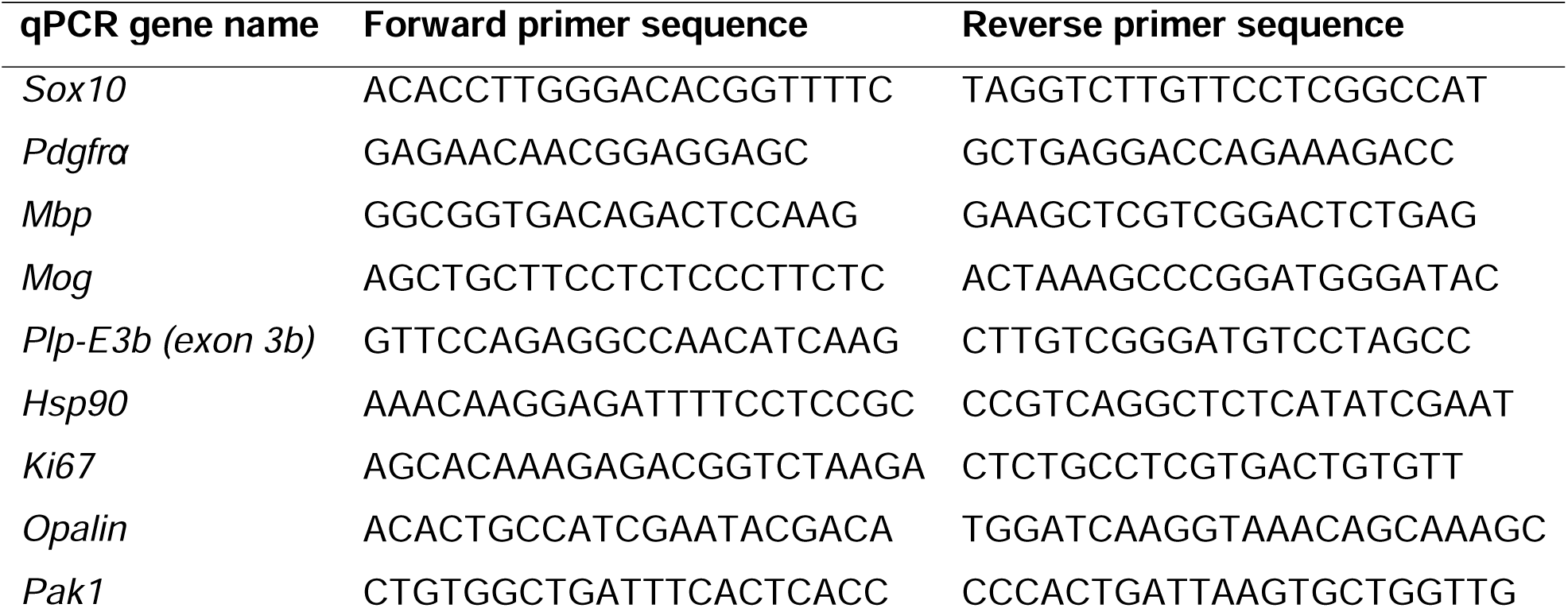

#### Tissue preparation and immunohistochemistry (IHC)

Mice were anesthetized using a ketamine/xylazine mixture and subsequently perfused transcardially with ice-cold phosphate-buffered saline (PBS). Tissues were harvested and immediately placed on dry ice for subsequent protein or RNA extraction, or fixed in fresh 4% paraformaldehyde (PFA, Electron Microscopy Science) for histological analysis. Tissues were post-fixed in 4% PFA for 2 hours at room temperature (RT), then washed with PBS three times for 15 minutes each. For cryopreservation, tissues were soaked in 30% sucrose (Fisher Chemical) in PBS overnight at 4°C before being embedded in O.C.T. compound (VWR International). Serial coronal sections (12 µm thick) were prepared using a Leica Cryostat (CM 1900-3-1) and stored at -80°C. For IHC, sections were air-dried at RT for 2 hours and blocked with 10% donkey serum in 0.1% Triton X-100/PBS (v/v) for 1 hour at RT. This was followed by overnight incubation with primary antibodies at 4°C. After washing with PBS containing 0.1% Tween-20 (PBST, v/v), sections were incubated with fluorescence-conjugated secondary antibodies for 2 hours at RT. DAPI was used for nuclear staining. Images were captured using a Nikon A1 confocal microscope. Sections with a 10 µm optical thickness were imaged via confocal z-stacking (step size: 1 µm) and compiled into a single, flattened image for quantification. Primary antibodies used were: PAK1 (1:100, #2602, Cell Signaling Technology), P-PAK1(Ser144) (1:100, #2606, Cell Signaling Technology), PDGFRα (1:100, #AF1062, NOVUS), MBP (1:200, #NB600-717, Novus), Sox10 (1:200, #ab155279, Abcam), Ki67 (1:100, #9129, Cell Signaling Technology), cleaved caspase3 (1:100, #9661, Cell Signaling Technology), PH3 (1:100, #9701, Cell Signaling Technology) and GFP antibody (1:200, #06-896, Millipore). Alexa Fluor®-conjugated secondary antibodies (1:500) are all from Jackson Immuno Research Laboratories.

#### Lysolecithin demyelination model and Neutral Red (NR) Labeling

Lysolecithin (LPC, Sigma) was administered into the corpus callosum of mice using a stereotaxic apparatus. Prior to the procedure, mice were administered buprenorphine (0.1 mg/kg, subcutaneously) for pain relief and anesthetized with a mixture of ketamine (0.1 mg/g) and xylazine (0.01mg/g). The injection site was precisely located at coordinates at +1.000 mm anterior and +1.000 mm lateral to the Bregma. The skull was carefully thinned with a drill, and a needle was inserted to a depth of 1.8 mm to target the rostral part of the corpus callosum. Using a microinjection pump, 2 μL of 1% LPC in 0.9% NaCl solution was injected over a period of 5 minutes. As a control, some mice received an injection of saline solution only. After injection, the needle was left in place for an additional 5 minutes to prevent backflow, then gradually withdrawn. P60 *Pdgfr*α*-CreER*^T2^:*Pak1*^fl/fl^ or *Pdgfr*α*-CreER*^T2^:*PID* and *Pdgfr*α*-CreER*^T2^ control mice were administered with tamoxifen followed by stereotaxic injection of lysolecithin into the corpus callosum 2 weeks after. Post-surgery, mice were placed on a heated water pad for recovery until they were ready to be returned to their home cages. Five days post-lesion (dpl), mice received an intraperitoneal (i.p.) injection of 500 μL of 1% Neutral Red (NR, Sigma-Aldrich) dissolved in PBS to label brain lesions. Two hours later, they were perfused intracardially with PBS, followed by the collection and fixation of brain tissue for histological analysis.

#### Immunocytochemistry (ICC)

Cells cultured on glass coverslips were fixed using 4% paraformaldehyde (PFA) for 30 minutes, then permeabilized with 0.1% Triton X-100 in PBS. Following permeabilization, cells were blocked with 10% donkey serum to prevent nonspecific antibody binding. After blocking, the cells were washed with PBS and incubated with primary antibodies overnight at 4°C. Subsequently, cells were treated with fluorescence-conjugated secondary antibodies at a dilution of 1:200 for 2 hours at room temperature. Nuclei staining was achieved using DAPI to facilitate visualization of cell nuclei. Fluorescent imaging was conducted with a Nikon A1 confocal microscope.

#### Plasmid transfection

Primary OPCs, at a density of 2 × 10^6, were seeded into poly-D-lysine (PDL)-coated wells of a 6-well plate one day before undergoing transfection. The transfection process utilized 6 μl of FuGENE (Roche Applied Science) and 2 μg of various plasmids, following the manufacturer’s guidelines. Specifically, cells were transfected with either 2 μg of an empty vector, 2 μg of a constitutively active PAK1 plasmid (pCMV6M-PAK1 T423E, Addgene plasmid # 12208 by Jonathan Chernoff), or 2 μg of a dominant-negative PAK1 plasmid (pCMV6M-PAK1 H83L H86L K299R, Addgene plasmid # 26592 by Jonathan Chernoff). This transfection was maintained for two days, after which the cells were processed for immunocytochemistry (ICC) and Western blotting assays.

#### Pak1 inhibitor treatment

For in vitro study, WT OPCs were seeded at a density of 10,000 cells per well on PDL-coated cover glasses within 24-well plates. One day post-seeding, the cells were treated with either the Pak inhibitor FRAX486 (15 nM), IPA3 (1 μM), or a vehicle control, all diluted in CGM medium. This treatment was maintained for three days. On the fourth day, 10 µM EdU was added to the CGM medium, and the cells were incubated for an additional 2 hours to facilitate EdU incorporation before the cells were fixed and permeabilized for subsequent analysis. The cells were cultured at 37°C in a humidified atmosphere containing 5% CO2.

For in vivo study, WT mice received intraperitoneal injections of IPA3 at a dose of 5 mg/kg/day or a vehicle control on postnatal days 2 (P2), 3 (P3), and 4 (P4). The mice were then prepared for analysis on P5. To label dividing cells, a 50 mg/kg dose of EdU was administered intraperitoneally 2 hours prior to tissue collection for histological examination.

#### EdU-labeling study

A 5-ethynyl-uridine (EdU)-labeling experiment was conducted using the Click-iT EdU Imaging Kits by Invitrogen, adhering to the provided manufacturer’s guidelines. Mice underwent euthanasia 2 hours post-intraperitoneal injection of EdU, dosed at 1 mg per mouse. Following euthanasia, brain specimens were harvested, fixed, and cryosectioned as described in the Tissue Collection and IHC protocol. The detection of EdU-labeled cells was achieved through the application of Click-iT reaction cocktails, also supplied by Invitrogen. Subsequently, the sections were incubated with primary and secondary PDGFRα antibodies and DAPI, before being mounted in the VectorShield mounting medium from Vector Laboratories. The mounted sections were then examined with an Olympus BX51 fluorescence microscope.

#### Magnetic activated cell sorting (MACS)

MACS was performed according to our published protocols (Zhang et al., 2021; Zhu et al., 2024). Whole mouse brains were dissected and cut into small pieces, then digested using the Neural Tissue Dissociation Kit (P) (Miltenyi Biotech), following the manufacturer’s guidelines. The resulting cell suspension was passed through a cell strainer to achieve a single-cell suspension. For the magnetic labeling of microglia, cells were incubated with anti-CD11b microbeads for 15 minutes at 4°C. Similarly, for the labeling of immature oligodendrocytes, the cells were treated with anti-O4 microbeads, and for astrocytes, with anti-ACSA2 microbeads, under the same conditions. Following each incubation with microbeads, magnetic separation was conducted using MS columns and the Octomacs manual separator. The isolated cells were flushed into tubes in 0.5% BSA in PBS (pH 7.2). After centrifugation, the cells were prepared for RNA extraction.

#### Quantification and statistical analysis

Data quantification was carried out by evaluators who were unaware of the genotypes and treatments. The results are presented as the mean ± SEM throughout this study. To visualize the data, scatter dot plots were employed, where each dot signifies an individual mouse or a separate experimental run. The Shapiro-Wilk test assessed the normality of the data distribution. For comparing two datasets, an unpaired two-tailed Student’s t-test was applied, with the degrees of freedom (df) indicated as t(df) in the figure legends. For analyses involving three or more groups, a one-way ANOVA with Tukey’s post-hoc test was utilized. The F ratio and degrees of freedom (numerator and denominator) were noted as F(DFn, DFd) in the figure legends, where DFn and DFd represent the degrees of freedom of the numerator and denominator, respectively. All graphical representations and statistical analyses were conducted using GraphPad Prism version 8.0. A P value of less than 0.05 was deemed statistically significant. The notation “ns” indicates a non-significant result, where the P value was greater than 0.05.

## Supporting information

Statistical analysis and parameters

PAK1's interactomes in OPCs and OLs identified by LC-MS/MS

GO terms of OPC interactomes of PAK1

GO terms of OL interactomes of PAK1

## Acknowledgements

We thank the funding agencies of NIH (R21NS125464, R01NS123080, R01NS123165, R01NS134887) and Shriners Hospitals for Children (85101-NCA-22, 85113-NCA-23 to FG, 85410-NCA-24 to YW). The authors declared no potential conflict of interest and all consented to publication.

