## Supplementary Figures for "Control of OPC proliferation and repopulation by the intellectual disability gene PAK1 under homeostatic and demyelinating conditions"

**Supplementary Figure 1 – mRNA expression of group I PAKs in oligodendroglial lineage cells quantified by scRNA-seq and qPCR (related to Figure 1)**

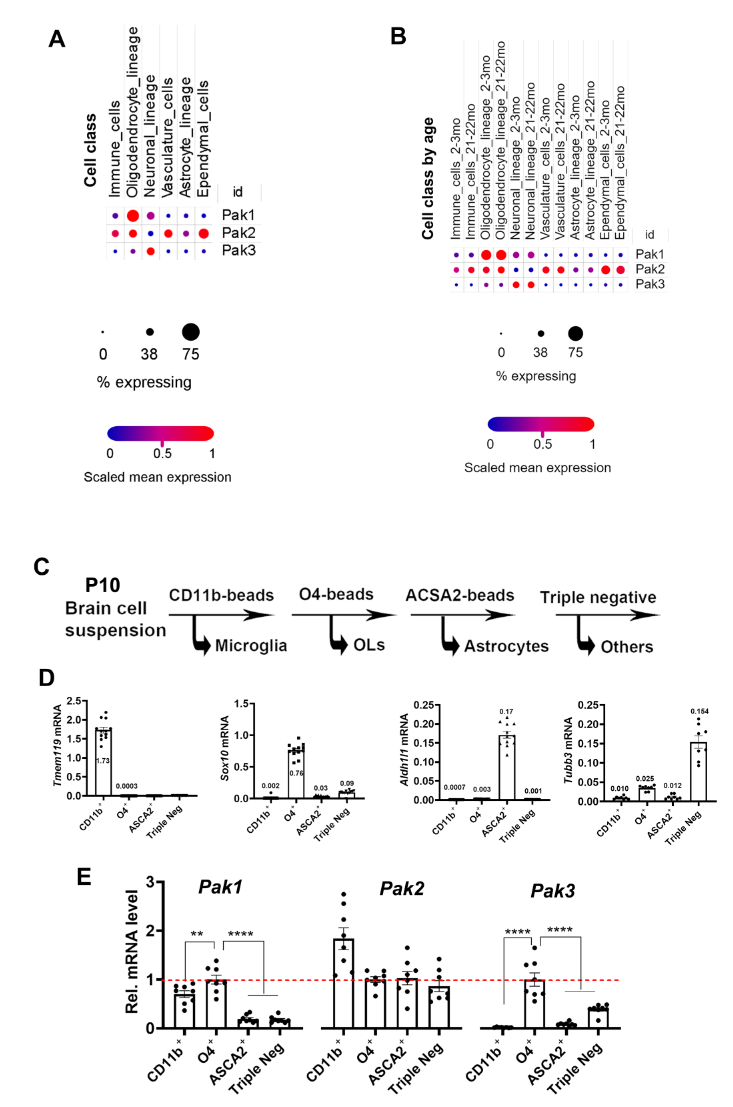

**A-B**, scRNA sequencing data at https://singlecell.broadinstitute.org/single_cell/study/SCP263/aging-mouse-brain (Ximerakis et al., 2019 Nat Neurosci - Single-cell transcriptomic profiling of the aging mouse brain showing group I PAKs expression at the single cell levels). **A**, heatmap plot of PAK1-3 mRNA expression by cell class. **B**, heatmap plot of PAK1-3 mRNA expression by cell class and ages of 2-3 months (young) and 21-22 months (aged) old mouse brain.

**C**, flow chart of acute isolation of different cell populations by MACS.

**D**, purity check by qPCR quantification of lineage-specific markers gene expression: *Tmem119* for microglia, *Sox10* for oligodendroglia, *Aldh1l1* for astroglia, *Tubb3* for neurons.

**E**, qPCR quantification of relative mRNA levels of Pak1, Pak2, and Pak3 in each cell component. mRNA expression levels were normalized to *Hsp90*. The expression level of PAKs1-3 was normalized to 1 in oligodendroglia (red dash line).

**Supplementary Figure 2 – (related to Figure 1)**

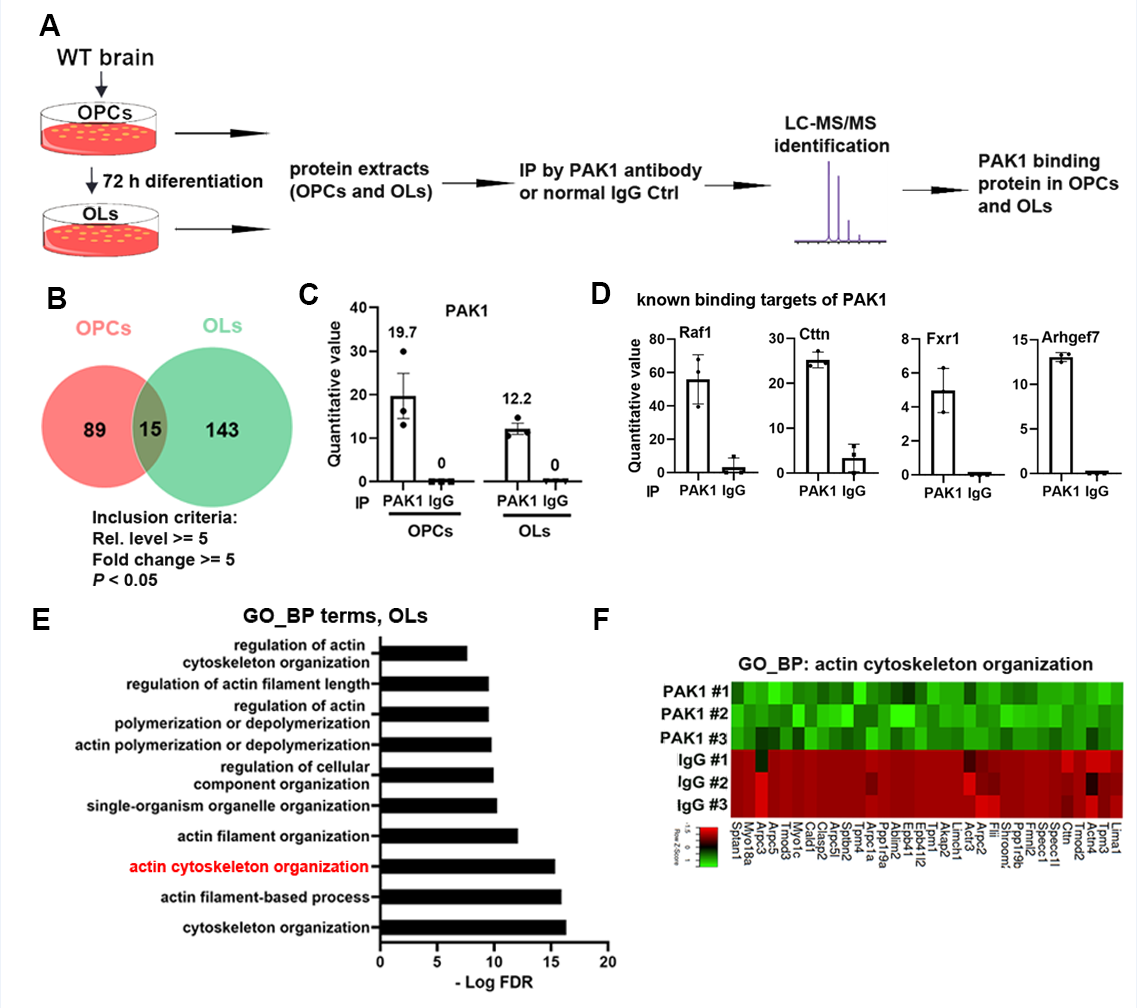

**A**, experimental designs for profiling PAK1’s interactomes in OPCs and differentiating OLs by proteomic approaches. IP, immunoprecipitation. LC-MS/MS, liquid chromatography with tandem mass spectrometry.

**B**, Venn gram showing the number of PAK1 binding partners in OPCs and OLs. Note that PAK1’s binding partners were largely non-overlapped in OPCs and OLs; only 15 proteins were identified both in OPCs and OLs. **C**, PAK1 was significantly enriched in anti-PAK1 IP component compared to IgG control component.

**D**, examples of known binding partners reported by previous studies.

**E**, GO_BP terms demonstrating significant enrichment of actin cytoskeleton organization among PAK1’s binding partners unique in OLs.

**F**, heatmap of identified binding partners of PAK1 in OLs that play a crucial role in actin cytoskeleton organization.

**Supplemental Figure 3 – inhibiting PAK activity is permissive for OPC differentiation into oligodendrocytes in the differentiation medium (related to Fig. 2)**.

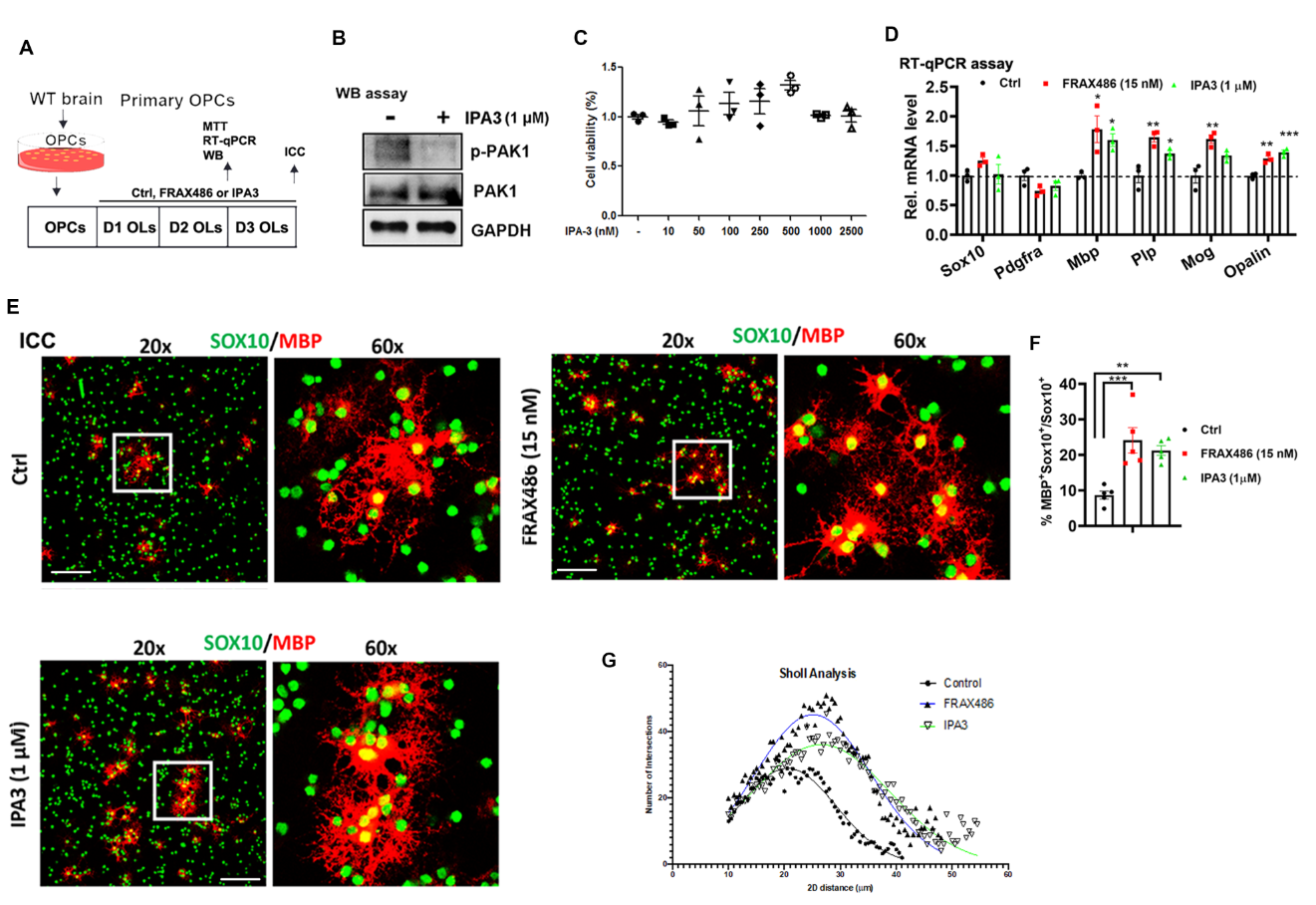

**A**, experimental designs of rat OPC differentiation cultured in the differentiation medium.

**B**, WB assay of p-PAK1 (Thr423) and total PAK1 at day 2 differentiation.

**C**, MTT cell viability assay of differentiating OL at day 2 in the presence of various IPA3 concentrations in the differentiation medium.

**D**, real time quantitative PCR (RT-qPCR) assay of *Sox10, Pdgfra*, and myelin gene *Mbp, Plp, Mog*, and *Opalin* at day 2 differentiation

**E**, ICC of Sox10 and MBP at low power (20x) and high power (60x) resolutions of at day 3 differentiation. Scale bar=100 µm.

**F**, quantification of percentage of MBP^+^Sox10^+^ cells among total Sox10^+^ cells at day 2 differentiation

**G**, Sholl analysis showing the number of intersections of MBP^+^ processes at various distances (um) from the center of OL cell bodies.

**Suppl Fig 4 - Strategy of PAK1 floxed mice and PAK1 deletion detection (related to Figure 3)**

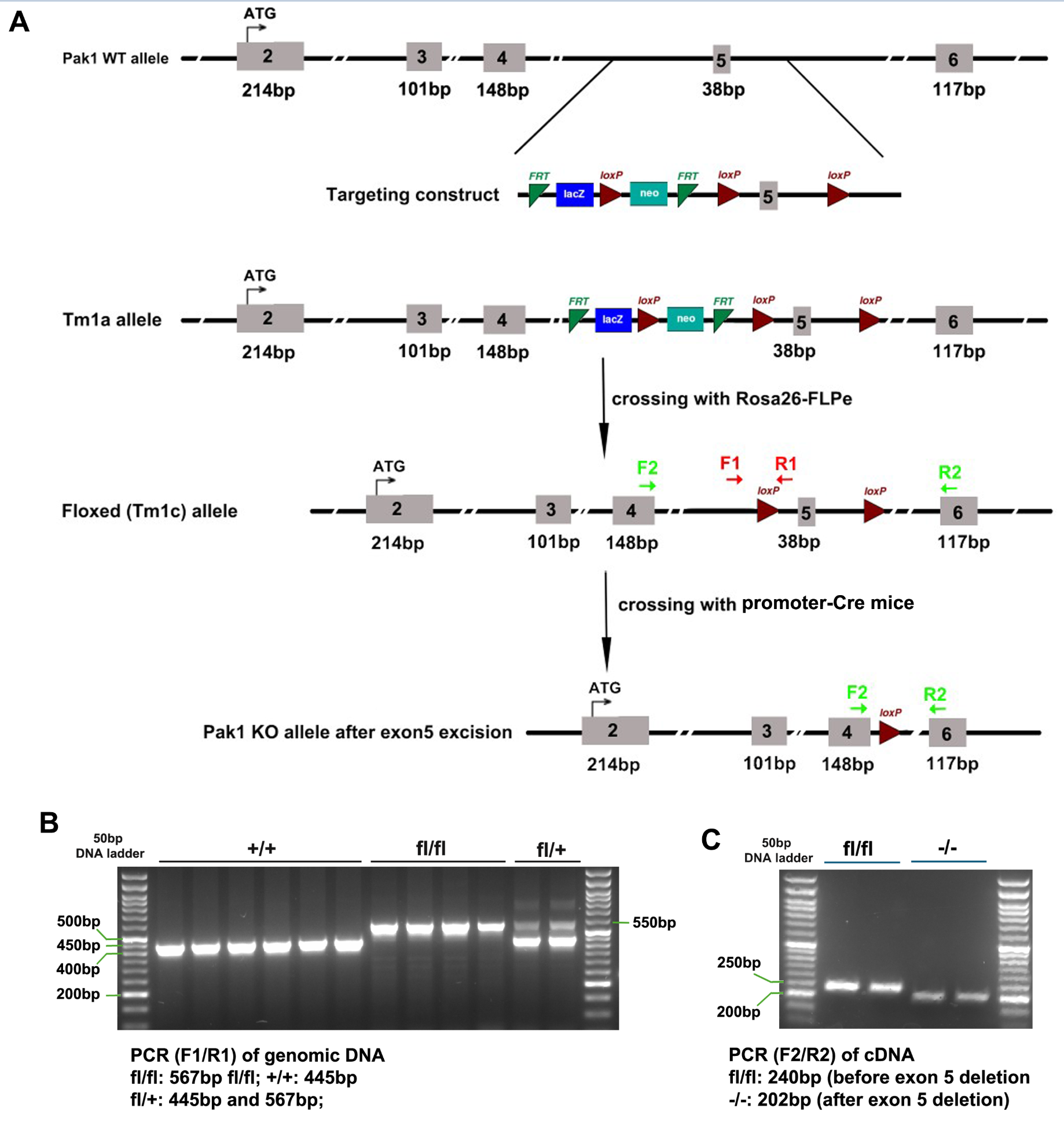

**A**, targeting strategy of Pak1-floxed transgenic line generation. The two loxP sites were inserted flanking the 38-bp exon 5. The primer pair of F1 and R1 were designed to detect Pak1 floxed alleles (Tm1c, fl/fl) using genomic DNA. The primer pair of F2 and R2 were designed to detect Pak1 exon 5 deletion alleles (-/-) using cDNA reversely transcribed from mRNA in Cre-positive cells.

**B**, gel images showing correct genotype results using genomic DNA. +/+, wild type alleles (445 bp), fl/fl, floxed alleles (567 bp), fl/+, heterozygous for floxed allele and wild type allele.

**C**, gel images showing correct results of exon 5 deletion using cDNA prepared from purified brain oligodendroglial cells by magnetic-assisted cell sorting (MACS of O4 antibody-conjugated magnetic microbeads). The brains were from P7 mice carrying *Pak1*^fl/fl^ mice and *Sox10-Cre:Pak1*^fl/fl^, respectively.

**Suppl Fig 5 – PAK1 inhibition reduces OPC proliferation and population expansion in vivo (related to Figure 3)**

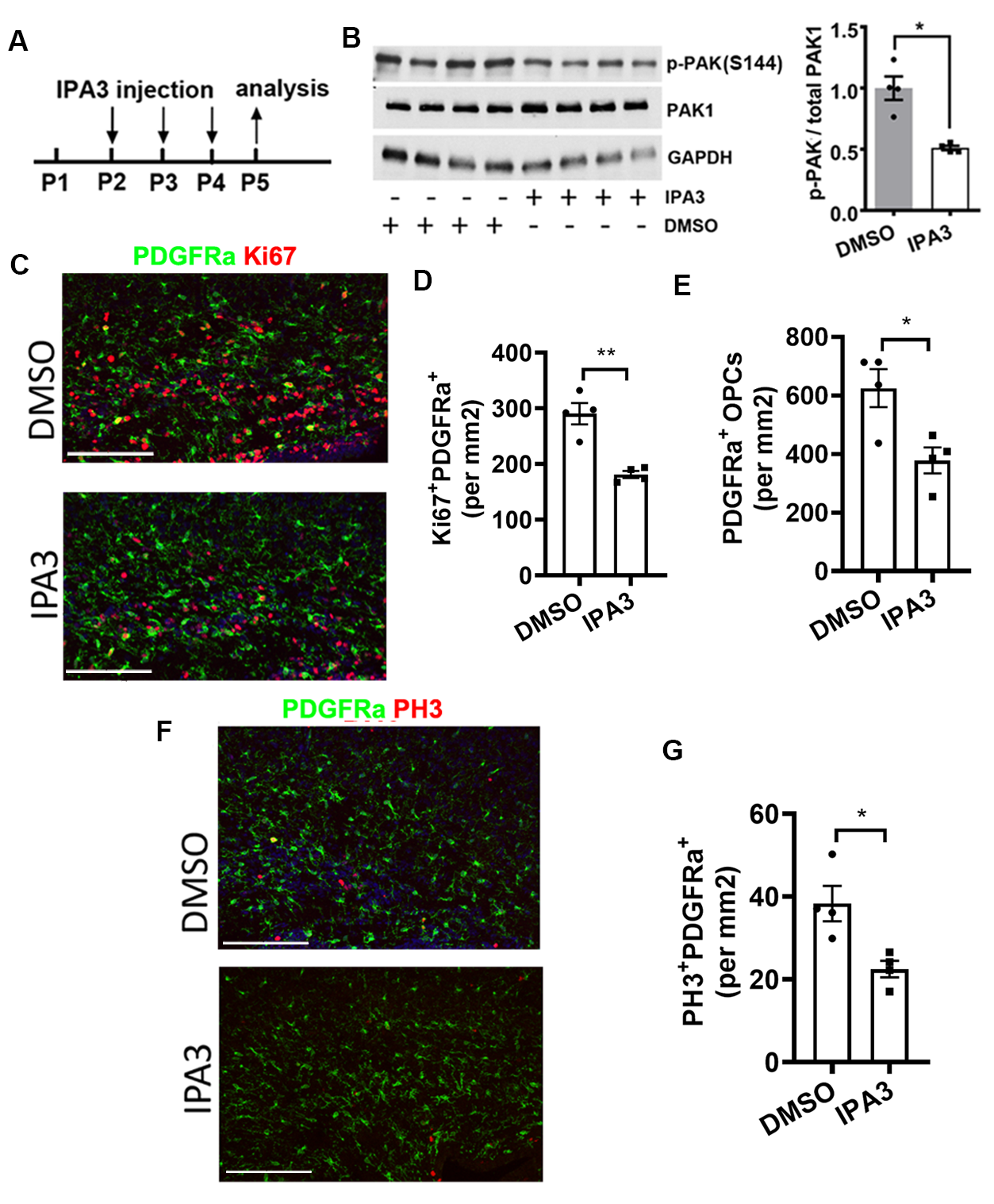

**A**, experimental designs for panel B-G. C57BL/6J wild type mice were i.p. injected with IPA3 at P2, P3, and P4 and analyzed at P5.

**B**, WB assay of p-PAK1 (Ser144) and total PAK1 in the brain. GAPDH was used as a protein loading control.

**C-E**, representative confocal images (**C**) and densities of Ki67^+^PDGFRa^+^ proliferating OPCs (**D**) and total PDGFRa^+^ OPCs (**E**) in the brain SCWM.

**F-G**, representative confocal images (**F**) and densities of PH3^+^PDGFRa^+^ mitotic OPCs in the brain SCWM.

Scale bars=100 µm.
